# SEMFA: A General Framework for Inferring Statistical Significance of Mahalanobis Similarity between Multi-Omics Profiled Samples Built on Multiple Factor Analysis

**DOI:** 10.64898/2026.06.18.733287

**Authors:** Jiali Han, Wenting Luo, Edwin Baldwin, Hao Helen Zhang, Lingling An, Jian Liu, Haiquan Li

## Abstract

**Motivation:** With rapid advances in sequencing technologies, many heterogeneous omics datasets have been generated, as seen in the Encyclopedia of DNA Elements (ENCODE) and many single-cell multi-omics sequencing projects, bringing substantial challenges to existing integrative methods. In this article, we report a novel multi-omics fusion and analysis software SEMFA which performs general parametric tests for the Mahalanobis **S**imilarity of samples based on the factor scores generated by an **E**xtended version of conventional **M**ultiple **F**actor **A**nalysis.

**Results:** Our developed method is effective and robust under both Gaussian and non-Gaussian assumptions. The mean F_1_ scores are over 0.8 when the column similarity level is 0.9 and the noise level ranges between 0.1 and 0.2, using simulation studies based on ENCODE count data. It was also efficient and effective at handling large-scale single-cell multi-omics data, as demonstrated in colon cancer cases as it unveiled signature network organization patterns of cells for stages III and IV.

## 1. Introduction

Driven by fast-advancing sequencing technology in the last decades, such as RNA-seq, ATAC-seq, CUT&Tag, and spatial sequencing, it is now a common practice to assay a sample from multi-omics scales, profiling them from genetic (e.g., copy number variation), transcriptomic, epigenomic, proteomic, and spatial perspectives, whether the samples are assayed in bulk or as single cells. For instance, the Encyclopedia of DNA Elements (ENCODE) (Harrow, et al., 2012), Roadmap Epigenomics Project (Bernstein, et al., 2010), and The Cancer Genome Atlas Program (TCGA) (Tomczak, et al., 2015) profiled many multi-omics assays for the same sample with bulk cells. Recently, more and more single-cell multi-omics profiling techniques have been developed for a variety of tissue types (Huo, et al., 2021; Ma, et al., 2020), especially for cancers (Nam, et al., 2021). Examples include scG&T-seq (single cell Genome & Transcriptome sequencing), scMT-seq (single cell Methylome and Transcriptome sequencing), scM&T-seq (single cell Methylome & Transcriptome sequencing), scTrio-seq (single-cell triple omics sequencing), and scCOOL-seq (single cell Chromatin Overall Omic-scale Landscape Sequencing) (Hu, et al., 2018). Multi-omics profiling provides more accurate assays for matched samples, reducing the effects of confounders, such as sample heterogeneity in multi-omics assays arising from different cohorts of samples with distinct omics groups. Control of sample heterogeneity facilitates the discovery of true mechanisms. With numerous applications, multi-omics profiling has already demonstrated its capability to unveil how genetic variations (e.g., single-nucleotide polymorphisms or SNPs) propagate through different levels of biology from DNA to the observable phenotypes. This includes how different biological entities (e.g., genes, proteins, regulatory elements, etc.) interact within and across scales, cell interactions with each other and pathogens, cellular organization, heterogeneity, and cell niches.

Amid the unprecedented opportunities created by multi-omics technology, they also bring a substantial challenge to integrate data from both computational and statistical perspectives. Early integrative methods for genomic profiles focused on integrating heterogenous multi-omics data that measured different samples with distinct omics, such as Seurat (Hao, et al., 2021). For multi-omics data measured on matched samples, fusion methods are preferred. As reviewed by us (Baldwin, et al., 2020) and others (Ballard, et al., 2024; Liu, et al., 2025; Ritchie, et al., 2015; Stahlschmidt, et al., 2022), fusion methods can be categorized by data fusion, model fusion, and mixture fusion which combines data and models within a single model. Data fusion, such as multiblock matrix factorization (Greene and Cunningham, 2009), multi-omics factor analysis (MOFA) (Argelaguet, et al., 2018), and other methods for multi-omic integration of bulk data including similarity network fusion (Wang, et al., 2014), iCluster (Shen, et al., 2009), and UINMF (Kriebel and Welch, 2022), are appealing as they can dramatically reduce the dimensions of large datasets while enabling various independent downstream analyses, such as clustering, classifications, regression, and mechanism discovery. Nevertheless, current data fusion methods (Bebek, et al., 2012) are still insufficient to address the challenges of multi-omics development because 1) existing fusion methods are not designed to handle vast volumes of data, both in terms of dimensionality (millions of features in genomic regions) and sample size (i.e., up to millions of single cells), generalized methods that are scalable to a large number of omics groups are still rare or costly to compute, few methods are designed to model the dispersed and sparse count data, including MOFA2 (Argelaguet, et al., 2020). Thus, further development of scalable, efficient data fusion methods is necessary to break the analytical bottleneck in multi-omics fusion-based analytics.

Further, a gap exists in the analytic space of the downstream analysis after dimension reduction of multi-omics data because very few methods have been developed to assess the statistical similarity of pairwise relationships. However, pinpointing these relationships can be essential in the biological world, such as cell-cell interactions, SNP-SNP interactions (Li, et al., 2016), and gene-enhancer interactions. For multi-omics data, few of them follow Gaussian distribution, but are generally in sparse and skewed zero-inflated binomial distributions (Zhang and Zhang, 2018), which further complicate the statistical inference of similarities.

In this article, we aim to address the knowledge gap in the multi-omics data fusion field. Our motivation is to use a straightforward, easy to calculate method to fuse multi-omics data rather than using sophisticated models as in the other methods. We will show that Multiple Factor Analysis (MFA) (Thurstone, 1931) is an appropriate framework for this purpose. It has been used frequently in other fields but insufficiently applied in bioinformatics (de Tayrac, et al., 2009; De Tayrac, et al., 2009; Voillet, et al., 2016). MFA can unveil simultaneous signals most in common across multi-omics scales, and project the samples into common scores in a low-dimension feature space across groups (e.g., omics groups). It can handle many omics scales and groups, even if a scale only has one single assay (one column). It is easy to interpret and has many intuitive properties in geometry space compared to competitor methods (e.g., autoencoders) (Ma and Zhang, 2019) because it is a linear transformation of the input data with a generalized singular value decomposition (Abdi, et al., 2013). Moreover, we will significantly extend the classic MFA framework to account for structure not only between the feature groups (e.g., omics types), but also between samples (e.g., samples of different individuals).

Even more importantly, we will propose a new framework to statistically assess the similarity between samples based on the most important factors with the major variance after MFA transformation. Generally speaking, there are three methods for constructing a similarity matrix: epsilon-neighborhood, k-nearest neighbor, or fully connected (Li, et al., 2021). Other methods include unsupervised clustering algorithms, such as Optimal Transport (OT) (Huizing, et al., 2022). As most omics profiles generate counting data, the assumption of Gaussian distribution may not hold for the factor scores. Compromised with the reality, we fit the factor scores with a proximate model and use the fitted parameters to estimate the statistical significance of the similarity. Using statistical theory and dynamic programming, we can calculate the probability and significance (e.g., p-value) within a reasonable time (seconds to minutes on a desktop computer) for large data even though the calculation involves multiple levels of integral (at least four levels). We also provide a general distribution as default fitting distribution, which is Asymmetric Laplace Distribution (ALD) (Koenker and Machado, 1999; Kotz, et al., 2001; Yu and Zhang, 2005), even our framework is open to any other distributions. For similarity measurement, we use Mahalanobis distance-based similarity because it counts for the variance in each dimension (Schissler, et al., 2015), which is uncorrelated between each dimension after MFA transformation. We named the software framework SEMFA, **S**imilarity based on **E**xtended **M**ultiple **F**actor **A**nalysis (SEMFA).

To demonstrate the effectiveness of the method, we will use simulated datasets and compare the feature reduction effects with state-of-the-art methods like MOFA2. More importantly, we examine the application potential of the method in knowledge discovery, using single-cell multi-omics data as an example. Thus, our contribution of the work will be extending the conventional MFA model, developing a new similarity test for counting data, and case studies of biological applications.

## 2 Methods

In this section, we will first describe how to conduct extended multiple factor analysis. Then, we will elaborate on how to prioritize pairwise similarity based on SEMFA transformed factor scores from input data following Gaussian distribution or any type of distribution (general cases). Finally, we will provide direction on SEMFA to generate simulation datasets and preprocessing of real datasets.

### 2.1 Data fusion of multi-omics Data by SEMFA

Suppose we have a data matrix (***X***; bold capital letter for a matrix) consisting of *n* rows of samples (or observations) and *m* columns for features generated from *g* groups (e.g., individuals) of samples and *k* types (or modals) of features (e.g., ChIP-seq arrays), as denoted in Eq. 1.

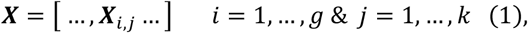

where ***X****i,j* where is the sub-matrices for group *i* and feature type *j*. Of note, the data matrix may be normalized, such as centered and/or scaled, prior to the analysis for meaningful biological implications.

Our framework utilizes generalized singular value decomposition (GSVD) (Abdi, 2007) to decompose the data matrix into singular matrices (***P, Δ***, and ***Q***), as shown in Eq. 2.

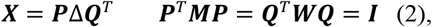

where ***P, Q*** are the left and right singular matrix of ***X, respectively;*** each column of ***P*** or ***Q*** is a singular vector. Further, **Δ** is the diagonal matrix of singular values of ***X*** with a dimension *r* at most the minimum of n and m. ***I*** is an identity matrix, and *T* is the transpose operator. If ***M*** and ***W*** are identity matrices, GSVD becomes singular value decomposition (SVD).

In MFA (Hotelling, 1933), the weights associated with each column group (e.g., assay) and row group (e.g., sample) are stored in diagonal matrices (***M*** or ***W***) and are used to weight the column and row groups, respectively. SEMFA employed the MFA framework to include groups of rows/samples to avoid dominance by a single group. By solving the GSVD for the data matrix, the decomposition results in common factor scores (***F***) of each sample on the subspace of representative features. The subspace, represented by the loadings (***Q***) of the right singular matrix can be interpreted geometrically as the common space shared among all features (Eq. 3).

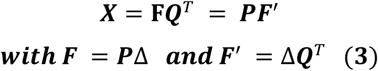

Similarly, each column can be projected as common factor values (***F***^’^) across various samples groups in the space of representative samples (***P***). The framework enables the extraction of concordant signals across multi-omics data, considering the structure of observations and variables (Abdi, et al., 2013).

In classic MFA, the weight of a sub-group, such as the weight for the corresponding sub-matrix (***X***_.*j*_ or simply ***X***_*j*_), is the inverse of the maximal eigenvalue of the sub-matrix, or the square of the first singular value (Abdi, et al., 2013). The weight is used for all columns for the sub-matrix.

Therefore, SEMFA could obtain concordant, common underlying signals across multi-scale data instead of being confined by the localized effects in each data group (e.g., omics) after extracting the essential information from the dataset, while counting for the observations and variables’ structure (Abdi, et al., 2013).

### 2.2 Measuring Similarity between Sample Pairs in SEMFA

We will apply Mahalanobis similarity/distance to measure the similarity (e.g., epigenomic) of sample pairs (*x*_1_ & *x*_2_) from the major factors of the input data ***X***. The Mahalanobis similarities count the variance in each factor to avoid any single factor’s domination and standardize the scores between 0 and 1. We employ a parametric estimation approach to evaluate the statistical significance of the similarity score and divide the original data distribution of **X** into two categories, which are Gaussian and non-Gaussian (e.g., zero-inflated negative binomial distribution).

#### 2.2.1 Mahalanobis Distance

Mahalanobis Distance (MD) quantifies the distance of sample pairs and further considers the covariance between dimensions of the data. The squared Mahalanobis distance (*u*) of two multi-omics samples (*x*_*1*_ and *x*_*2*_) is defined as the sum of the squared Mahalanobis distance on each projected dimension after MFA,

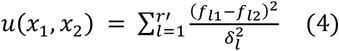

Where *f*_*l1*_ and *f*_*l2*_ are the factor scores of the sample *x*_*1*_ and *x*_*2*_ on the projected dimension *l* respectively, r^*’*^ is the number of dimensions considered, where *r*^’^≤*r*, the maximal number of singular values after GSVD, and *δ*_*l*_ is the estimated standard deviation of factor score in SEMFA dimensional *l*, which is proportional to the singular value in the dimension.

#### 2.2.2 Mahalanobis Similarity

Mahalanobis Similarity standardizes the similarity of each factor (*l*) by exponentiating the Mahalanobis distance with a negative coefficient of β to adjust the scale of the similarity due to the limitation of computing accuracy in computers. This normalizes the similarity measures to values between 0 and 1, corresponding to no similarity and identity respectively.

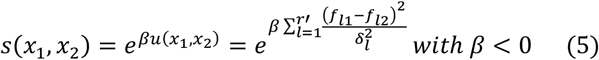

### 2.3 Mahalanobis-based Tests of SEMFA for Sample-similarity from Gaussian Distributed Genomic Data

#### 2.3.1 Parametric Test of Mahalanobis Distance

If we assume the original dataset is centered and follows a multivariate Gaussian distribution, the factor score distance (*y*_*l*_ *= f*_*l1*_ - *f*_*l2*_) on each dimension of the SEMFA space for a pair of samples (*x*_*1*_, *x*_*2*_) still follows a univariate Gaussian distribution with mean zero and variances 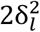. Then, the Mahalanobis distance of the two samples can be represented as a function of factor score distance in all dimensions,

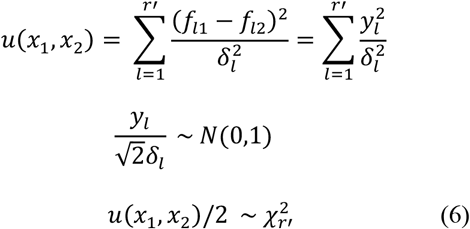

Thus, Mahalanobis distance follows a generalized chi-squared distribution with the degree of freedom as the number of SEMFA dimensions, *r’*. Specifically, it is a weighted (weight of 2) chi-squared distribution (last two lines of Eq. 6).

#### 2.3.2 Parametric Test of Mahalanobis Similarity

Since MD follows a generalized chi-squared distribution under the assumption that the input data follows Gaussian distribution by the above subsection, we can use the probability of transformations of the random variable distance (*u*) to calculate the similarity (*s*). Assuming the input data is centered after transformation, the distribution of a similarity value *s* is given by Eq 7,

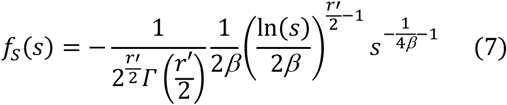

### 2.4 Mahalanobis-based Tests of SEMFA for Sample-similarity from Non-Gaussian Genomic Data

In general, the original dataset that is used to quantify the similarities does not follow a Gaussian distribution. Instead, studies revealed that sequencing-derived count data fitted better with zero-inflated negative binomial distribution or zero-inflated Poisson distribution (Cho, et al., 2020; Mesner, et al., 2013; Ren and Kuan, 2020). Here, we deploy a more generalized statistical model which passes over the Gaussian assumption and captures the natural over-dispersion of datasets in the ENCODE (Ren and Kuan, 2020) and single-cell sequencing data. After the linear transformation and combination of the SEMFA, the factor scores in each dimension become more complicated and may no longer follow a zero-inflated negative binomial distribution. Thus, we should consider any type of distribution for the factor scores in each dimension. For any such distribution, we can either use a probability density function (*f*) or the characteristic function (*φ*) (which is the Fourier transformation of the probability density function) to describe the distribution of factor scores, because there is a one-to-one correspondence between the two functions.

#### 2.4.1 Density Function of Mahalanobis Similarity of Sample Pairs from Data of Any Distribution

Again, assume *f*_*l1*_ and *f*_*l2*_ are two independent factor scores in the *l*-th dimension of a pair of samples (*x*_*1*_, *x*_*2*_), and *y*_*l*_ is the difference of them. Let φ_*f*1_ and φ_*y*1_ be the characteristic function of *f*_*l1*_*/f*_*l2*_ and *y*_*l*_, respectively. Using transformation theorem of random variables, the density function of Mahalanobis similarity of two samples *x*_*1*_ and *x*_*2*_ is given by

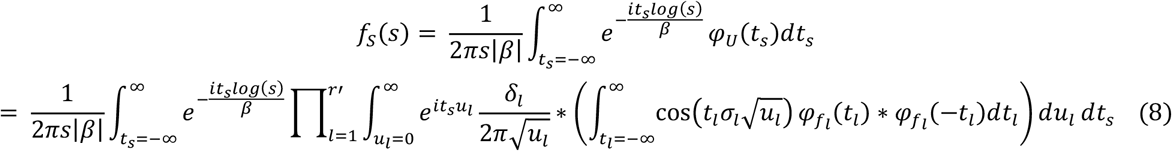

The probability function will be used to calculate the parametric p-values. For implementation, we have used dynamic programming to cache the intermediate results in each level of the integration with the hashmap package in R (Russell, 2017), so that the four levels of the integral can be calculated in a reasonable time.

#### 2.4.2 SEMFA Analysis of Sample-pair Similarity by Asymmetric Laplace Distribution based Approximation

We found that Asymmetric Laplace Distribution (ALD) fitted factor scores transformed well from zero-inflated count data and is a very flexible distribution that allows all numeric values. Therefore, we used ALD as an example for negative binomial sequencing data to simulate the factor scores on each major dimension. Generally, there are two types of definitions for the probability density function for ALD. In our study, we use the one modeled by *μ, σ*, and *p*, where *μ* is the location parameter, *σ* is a scale parameter and *p* is a skewness parameter (Jammalamadaka and Kozubowski, 2004; Kotz, et al., 2001).

Denote *f*_*l1*_ and *f*_*l2*_ as the factor scores of a pair of samples (*x*_*1*_, *x*_*2*_) on the dimension *l*, which approximately follows *ALD(μ*_*l*_, *σ*_*l*_, *p*_*l*_). The characteristic function of ALD is:

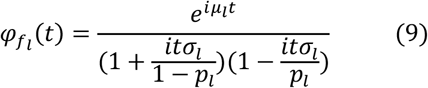

The probability density function of Mahalanobis Similarity *s* is shown in Eq. 10 if we replace the singular value *σ*_*l*_ with the estimated standard deviation since the estimated variance may be slightly different from the singular value:

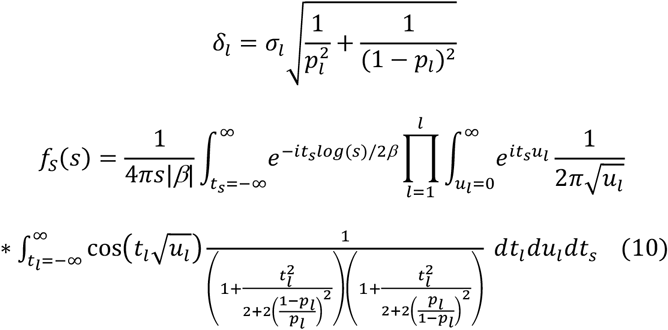

where *s* is similarity defined in Eq.5. Proof omitted.

#### 2.4.3 Mahalanobis Similarity Test Design with ALD

In the parametric method, we use the probability density function of Mahalanobis similarity to calculate the p-value for each sample pair on major factor scores. For example, the number of factors with accumulative variance is more than 80% of the total variance. By default, we use ALD fitting factor scores in each dimension, as in Eq. 10, but also support a customized version of characteristic function in Eq. 8 if the function of each dimension is provided by a user.

### 2.5 Simulation Method using ENCODE data for SEMFA

To evaluate our method, we used multi-omics data for one of the most studied lymphoblastoid cell lines, GM12878 in ENCODE, to study the fusion effect of multi-omics data. We downloaded the data from ENCODE on October 23, 2017 and processed their signals through VCFtools (Danecek, et al., 2011). Then, we mapped the signals to lead SNPs reported by the GWAS Catalog (MacArthur, et al., 2016) and their Linkage disequilibrium SNPs (LD; r^2^ > 0.8) based on LD results among SNPs in 1000 Genomes Project (Clarke, et al., 2012). We removed identified columns caused by different versions of the data and empty columns with no signals. Finally, the processed datasets left 48,337 SNPs and 315 columns from 12 types of assays, such as RNA-seq, ChIP-seq, DNase-seq, and RIP-CHIP, etc.

We estimated the mean and variance/covariance structure from the dataset and used that information for simulation. We randomly generated 100 SNPs using the mean values and correlation structure for the columns of a subset of groups (e.g., seven) where the count data was available. Then we generated signals buried in the randomly chosen SNP pairs (according to a percent (e.g., 1%) of total 4,950 pairs) considered to have high epigenomic similarity. We modified one of the SNPs (similar SNP) based on the other SNP (anchor SNP). We first set the proportion of columns (such as 80% of total columns, referred as similarity level) of the similar SNP to the value of the corresponding column of the anchor SNP, then added normally distributed noise with a proportion (e.g., 0.1) of the original standard deviation from each column.

For the simulation study based on Gaussian distribution, the dataset was generated by the multinormal distribution simulation mvrnorn function in the MASS package (Venables and Ripley, 2013). For zero-inflated non-Gaussian distribution data, we used multivariate zero-inflated negative binomial data generating function rmvzinegbin in the SpiecEasi R package (Kurtz, et al.).

SEMFA was applied to integrate these datasets and infer the statistical significance of the 4, 950 pairs among the 100 simulated SNPs (Methods 2.2 - 2.4). We used Gaussian-based factors and ALD fitted distribution for Gaussian and non-Gaussian data respectively. To evaluate the effectiveness of the parametric approach, F_1_ score, which is the harmonic mean of precision and recall, is calculated for comparison purposes. We exploited different parameters to examine the robustness of the method and compared the feature reduction effect by SEMFA with the state-of-the-art multi-omics integration method MOFA2 and naïve method of PCA.

### 2.6 Preprocessing of Single-Cell Multi-Omics Data of Human Colorectal Cancer

To test the effectiveness of the method for biological discovery and computational efficiency, we downloaded the large (zipped file size 717.9gb) human colorectal cancer data GSE97693 (Bian, et al., 2018). The dataset consists of over 1,000 cells from four regions of 10 patients with colon cancer stage III and IV, plus Hep-2 cell line as control cells. We chose 616 cells from six patients (CRC01, CRC02, CRC04, CRC09, CRC10, and CRC11) and the cell line for our study, as all the cells were measured by both RNA-seq and Bisulfite sequencing (scBS-seq) of the scTrio-seq2 on the same cell. The scRNA-seq reported 21,186 genes, while methylation sites from scBS-seq were summarized into 885,420 regions through automatic clustering that allowed up to 100 base pairs between adjacent sites in any cell. We conducted SEMFA analysis and grouped the columns by omics type (two groups) and grouped cells by patients. Then, we calculated the pairwise similarity with the parametric test using ALD fitting. After getting the p-value for each similarity, we adjusted them by false discovery rate (FDR). We plotted a network for the cells showing the three nearest neighbors with significant similarity (FDR<0.05) and colored the cells by regions and shaped them for patients, to examine whether the network provides novel insight into the underlying biology.

## 3 Results

### 3.1 Performance Evaluation of SEMFA under the Gaussian Assumption

We obtained 7 groups of assays with a total of 231 columns for the Gaussian-based simulation, which were ChIP-seq, DNase-seq, FAIR-seq, RIP-chip, RIP-seq, RRBS, and whole-genome shotgun bisulfite sequencing. We conducted SEMFA and compared the results with that of MOFA2 (with default parameters) using the 100 simulated datasets where a proportion of similar columns varied from 70% to 90%, noise levels varied from 10% to 20%, and the number of factors accounted for at least 80% of total variance in the similarity calculation. The coefficient *β* used in the similarity approach was set as -2. **Figure 1** shows the number of factors necessary to reach a total of 80% variance with a similarity level of 70% and noise level of 0.2 using the simulated datasets. Compared with PCA and MOFA2, SEMFA contains more information in the top factors when the features are not scaled. SEMFA had larger cumulative variance than PCA and MOFA2 given the same number of top factors. Other settings of simulations yielded similar trends when features were not scaled.

**Fig. 1.**
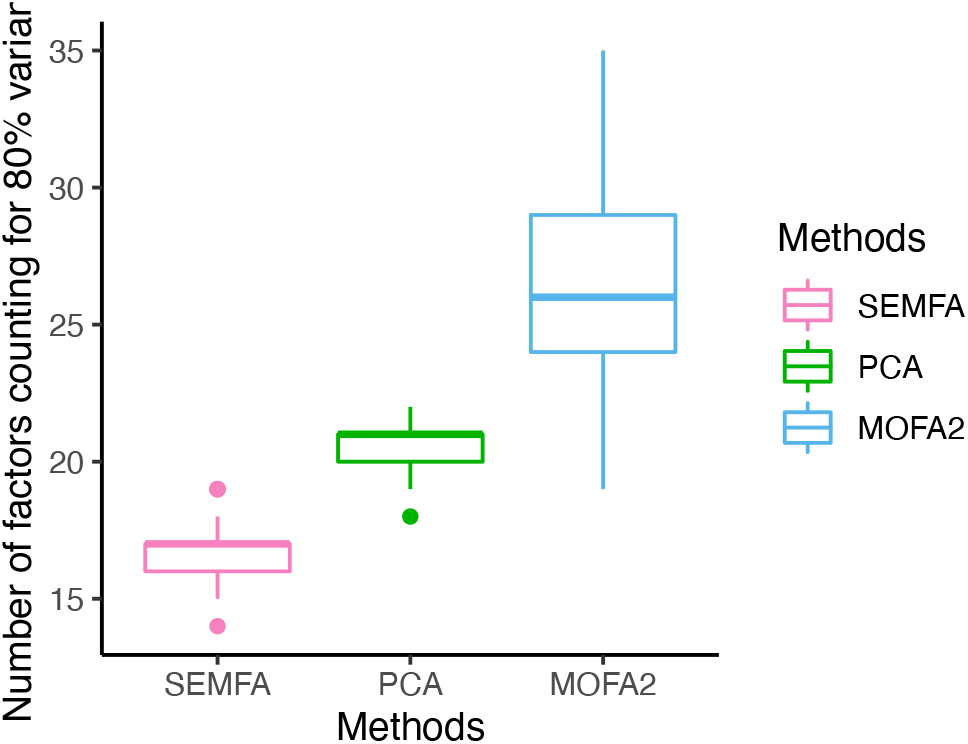
Number of top factors that reach 80% of the total variance of the dataset. Boxplot shows the number of factors contributing to at least 80% variance for SEMFA methods, PCA, and MOFA2 for 100 simulated datasets with 231 columns of data from 7 groups following Gaussian distributions. Coefficient β is set as -2, column similarity level is set as 0.7, and noise level is set as 0.2. Columns of data were centered but not scaled before feature reduction.

**Fig. 2** shows the distributions of maximal F_1_ scores from each of the 100 simulated studies with varied simulation parameters using SEMFA-based feature reduction and testing on standardized datasets. SEMFA achieved high F_1_ scores, with mean F_1_ scores ranging from 0.779 to 0.881 when the signals and noises varied in the specified range. The standard deviations of our F_1_ scores were 0.037 to 0.060, indicating the robustness of these approaches. The corresponding p-values achieved the max F_1_ score of each approach in each simulation iteration ranges from 0.0002 to 0.07, following convectional significance cutoff after multiplicity control.

**Fig. 2.**
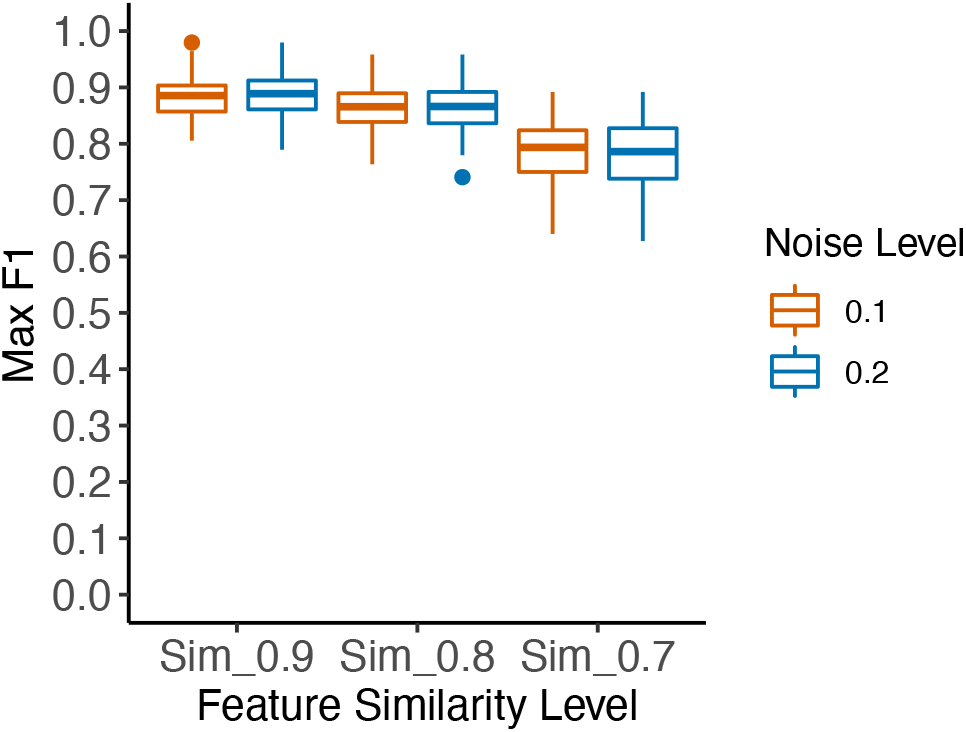
F_1_ Scores of SEMFA-based Mahalanobis Similarity (MS) test from Gaussian simulated data. Noise levels 0.1 and 0.2, respectively. Coefficient β is set as -2, column similarity levels are set in the range of 0.9, 0.8, 0.7. Columns of data were standardized prior to conducting SEMFA.

We exploited different settings for the coefficient *β* to examine the influence of the parameter on the performance of the proposed method. The coefficients only made a tiny difference when the absolute value exceeded a limit which made all similarities 0, going beyond the precision of computer float calculations (power of exponential less than -746).

### 3.2 Performance Evaluation of SEMFA under non-Gaussian Assumption

We used the same mean and correlation structure from the 7 groups of GM12878 processed data in section 3.1 to start the simulation of zero-inflated negative binomial data using an inflation ratio of 0.3. However, only 4 groups of simulated omics types were obtained after removing columns with all zeros. Omics groups with a few columns failed in generating any non-zero data. The distribution of F_1_ scores for each of the 100 simulated studies on standardized datasets is shown in **Fig. 3**, with mean values ranging from 0.585 to 0.83, while standard deviations range from 0.043 to 0.076. P-values reached the maximal F_1_ score ranging from 0.0003 to 0.002. Coefficient *β* was a little more sensitive due to the overdispersion of negative binomial data so the absolute number could not be too large (e.g., >100). The results demonstrate that the proposed method can be extended to non-Gaussian distributions. It should be noted that we failed to converge during model optimization using the only non-Gaussian model of Poisson in MOFA2, making a fair comparison of feature reduction unavailable. For PCA, the first factor always takes the majority of (>80%) variance due to the over- dispersed data whether it is scaled or not.

**Fig. 3.**
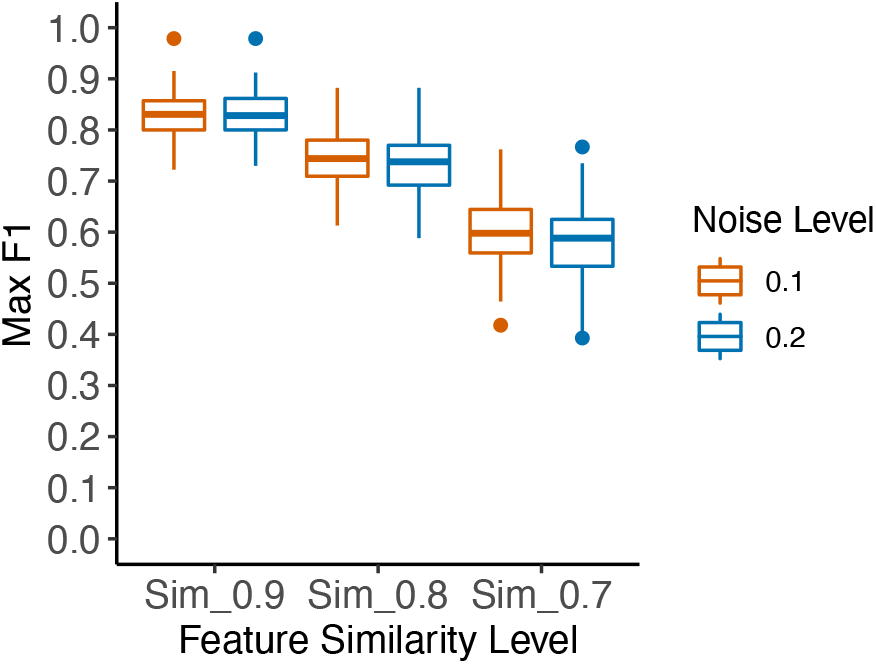
F_1_ Scores of parametric tests of Mahalanobis similarity under non-Gaussian Assumption. Coefficient β is set as -*2*, column similarity level is set as 0.9, 0.8, and 0.7, and noise level is set as 0.1 and 0.2. Columns of data were standardized prior to conducting SEMFA.

### 3.3 Performance Evaluation of SEMFA by single-cell data

To assess the performance of our proposed data fusion method based on SEMFA framework and Mahalanobis similarity, we preprocessed the GSE97693 dataset from GEO (see Methods) and tried to retain most of the raw signals before running SEMFA. We conducted both SEMFA and PCA/SVD analysis using the large dataset consisting of two groups and 906,606 features. We compared the first two factors by SEMFA and PCA of the concatenated data (no omics weight in PCA but both omics and patients are weighted by SEMFA; data are centered but not scaled before SEMFA/PCA) and on each of the individual omics scales (scRNA-seq and scBS-seq separately). We examined which feature reduction method can better recapitulate stages of colon cancer using single-cell data with cell-matched scTrio-seq2 assays using only top factors.

The dimension reduction results are shown on **Fig. 4**. It shows that SEMFA analysis of the whole datasets (Fig. 4D) achieved the best separation via clustering of colon cancer stage III (green), IV (blue), and control cells. Even though an individual scale (e.g., scBS-seq; Fig 4F) may not separate phenotypes well, the fused data (Fig. 4D) took the most convergent and relevant signals across scales. The fused data clustered the phenotype signals better than either of the individual signals while using only the first two factors (components), demonstrating the advantage of data fusion. More importantly, the factor scores of the fused data (Fig. 4D) and factor scores for each individual omics scale (called partial factor scores (Abdi, et al., 2013)), which are scRNA-seq (Fig. 4E) and scBS-seq data (Fig. 4F), correspond to the same SEMFA feature space. In fact, the former is the mean of the other two (the contribution of the omics to the factor scores, or partial factor scores with the same factor space). SEMFA analysis from the whole dataset outperformed PCA analysis (Fig. 4A) using concatenated data without weighting, as demonstrated by the separation of control cells. The control cells are located slightly outside of the cluster of stage IV cells in SEMFA analysis but mixed with stage IV cluster by PCA analysis. This occurs even though PCA analysis on the whole dataset yielded better separation of cancer stages than its analysis on individual scales (Figs. 4B × 4C).

**Fig. 4.**
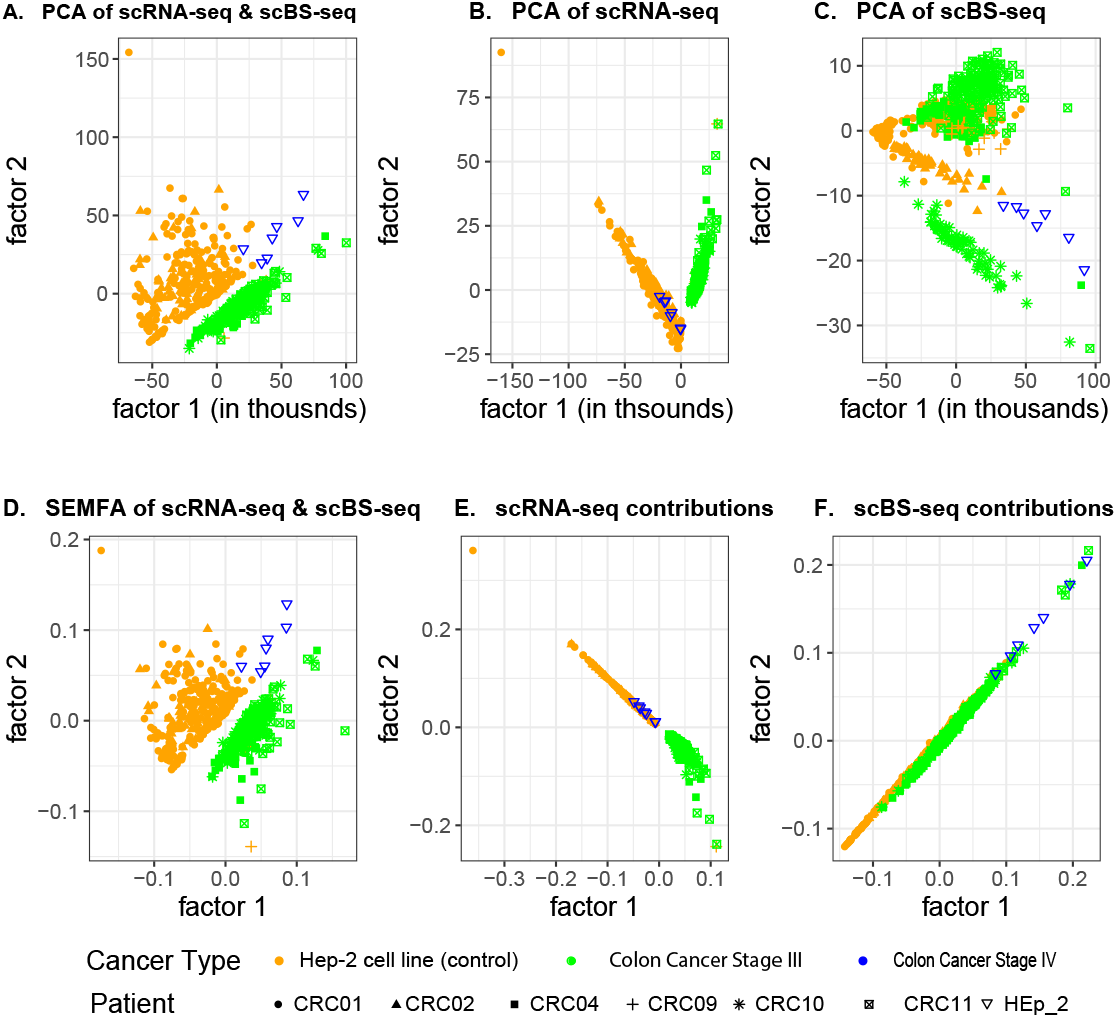
Data fusion with Multiple Factor Analysis (SEMFA, D-F) outperformed Principal Component Analysis (PCA, A-C) in single-scale data when recapitulating stages of colon cancer in the cell-matched scMultiOmics dataset GSE97693 using the first two factors.

Finally, we calculated the statistical significance among the 616 cells using our ALD-based test (Methods 2.4.2). We plotted the significantly similar cells (FDR<0.05) that are among the three-nearest neighbors of a cell within a network using a similarity test of the first 14 factors counting for over 80% of variance (**Fig. 5**), which resulted in 533 cells and 1,548 connections. The network shows the cell similarity or connection in terms of biological profiling. The three-nearest neighbor algorithm naturally connects cells of a patient together, indicated by six small clusters corresponding to each patient (coded by the shape of the node). More importantly, it unveiled an important pattern that separated cells of stage III (CRC01, CRC02, and CRC09) and IV patients (CRC04, CRC10, and CRC11). For stage III cells, primary tumor cells (in blue) tend to connect with each other, along with lymph node metastasis cells (in green). Two separate clusters can be easily seen from each stage III patient even though the two clusters may be connected. For stage IV patients, primary tumor cells are scattered within the network and connected more frequently with post-treatment liver metastasis cells (pink), liver metastasis cells (orange), and lymph node metastasis cells (green). Of note, only CRC01 (circle) had post-treatment cell sequencing. More interestingly, cells of distinct patients (CRC01 and CRC02) in the same stage (IV) connected with each other and formed a big cluster, except for CRC09, which only had a few cells pass quality control. This recapitulates the knowledge that late-stage patients have cells which are less differentiated and may distantly metastasize to other organs, leading to similar genomic profiles, even more than cells in adjacent spaces.

**Fig. 5.**
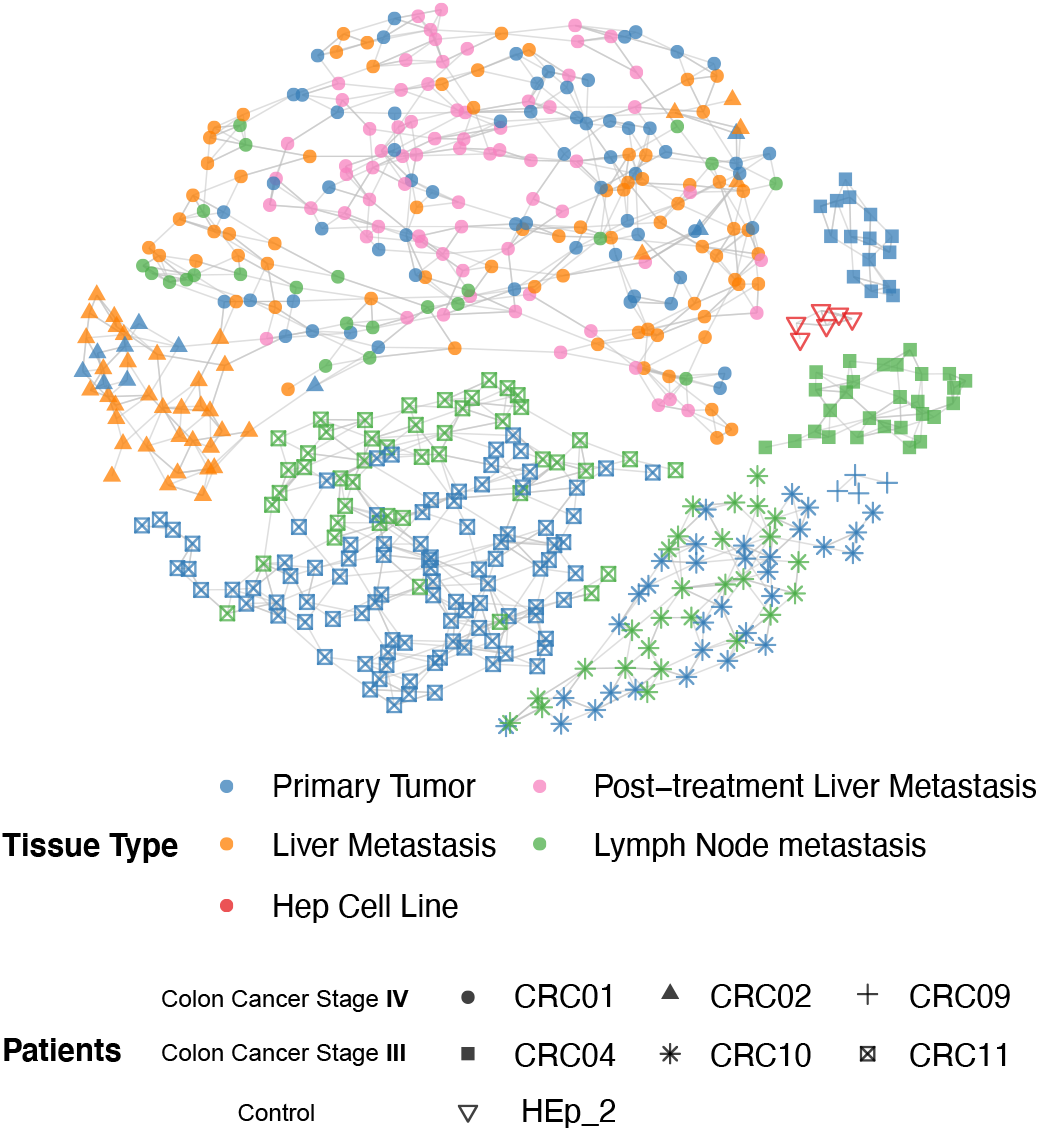
Plot of the tumour regions networks among the six patients. It visualizes the network where the colour and shape of nodes express the tumour regions and patients, respectively; lines between the nodes express significant similarity. The resulting networks exhibited the inner association of cells calculated by the similarity while the tumour stages are clearly defined with different clusters.

## 4 Conclusion

In this study, we reported a multi-omics framework SEMFA, that extended the conventional multiple factor analysis (MFA) to balance the effect from sample groups, such as a diverse number of cells from an individual in the colon cancer dataset GSE97693. SEMFA can handle a variety of samples (from a few to millions) containing a large number and varied size of feature groups, as we demonstrated with nearly a million. The feature reduction is nonparametric and thus much faster in calculations than other parametric ones like MOFA2. We have also shown that the SEMFA helped uncover verifiable signals (cancer stages and types of cancer cells). From both simulation and real data, we found that SEMFA is more efficient in feature reduction for non-centered Gaussian data and that the top factors kept a higher proportion of variance as compared to MOFA2, even though MOFA2 used sophisticated model inference. The highly reduced dimensions allowed critical phenotypic information to be distinguished, like cancer stage, compared to PCA (Fig. 4).

More importantly, we developed a novel statistical method to infer the significance of pairwise similarity which is based on reduced dimensions. We make no assumptions about the input data, having to determine and fit a distribution if the distribution of the input data is unknown. While the multi-level integral (at least 4 levels) is usually intangible in computation, we achieved a fast algorithm using a dynamic program. The default ALD distribution was found to be efficient at handling highly skewed data in a reasonable time (seconds to minutes for calculating p-values for thousands of similarities). We have demonstrated the accuracy of the similarity and robustness to noise levels (very little effect on the accuracy) using simulation data. We also demonstrated the capability to unveil single-cell relationships, providing novel insight on cancer-cell relationships and organizations.

Our similarity measure is very general. It can be used on data from many other sources other than transformed data from SEMFA. In the future, we will integrate MOFA2 into our similarity test framework. In this manuscript, we only reported similarity from samples, but our framework can be easily extended to map features into a representative sample space. With this, feature relationships can be measured similarly, but it is out of the scope of this paper and will need to be reported elsewhere. Of course, the framework will be limited to a manageable size of factors (up to hundreds) for similarity inference due to integral computing and may need a version with high-performance computing if the number of samples exceeds multiple millions, since the relationships cannot be stored in a typical computer’s memory.

In this work, we chose Mahalanobis similarity over Mahalanobis distance because it is easier for the integral to converge, plus its similarity to the chi-squared distribution. We chose Mahalanobis similarity over Euclidean-based similarity to count for the contribution of factors with less variance. We will keep studying which set of factors can better represent similarity among samples (features) as larger variance factors and smaller variance factors may have different biological meanings for multi-omics data.

The methodology has some limitations. 1) We assume independence among different factors which may be inaccurate since factor scores based on SVD can only guarantee uncorrelation but not independence. Nevertheless, we observed nearly zero dependence (absolute correlation<10^-16^) among different factors of colon cancer samples. 2) In rare cases, the integral may fail to converge. We will improve the adaptive integral algorithms to make them more flexible and robust on parameter settings. 3) The case study may be dominated by highly methylated regions where other preprocessing methods may be more appropriate, such as using the mean signal in a clustered region.

For future work, we will study more distance and similarity measures while testing more types of data with various distributions. We will compare more competitive tools, such as Seurat (Hao, et al., 2021), and extend or integrate our tools to work on heterogeneous sample integration.

In summary, the SEMFA provided a general and efficient tool that enables various downstream analyses, such as the clustering demonstrated in our case studies. The similarity test is statistically and computationally innovative, promising to lead to more applications, such as single-cell studies of cell-cell interactions, tracing of metastasis, or cellular organizations in three-dimensional space.

## Acknowledgements

We appreciate valuable comments from Dr. Edwin Baldwin.

